# Tracking the Subtle Mutations Thriving Host Sensing by the Plant Pathogen *Streptomyces scabies*

**DOI:** 10.1101/097998

**Authors:** Samuel Jourdan, Isolde M. Francis, Benoit Deflandre, Rosemary Loria, Sébastien Rigali

## Abstract

The acquisition of genetic material conferring the arsenal necessary for host virulence is a prerequisite on the path to become a plant pathogen. More subtle mutations are also required for perception of cues witnessing the presence of the target host and optimal conditions for colonization. The decision to activate the pathogenic lifestyle is not ‘taken lightly’ and involves efficient systems monitoring environmental conditions. But how can a pathogen timely trigger the expression of virulence genes if the main signal inducing its pathogenic behavior originates from cellulose, the most abundant polysaccharide on earth? This situation is encountered by *Streptomyces scabies* responsible for common scab disease on tuber and root crops. We here propose a series of hypotheses of how *S. scabies* could optimally distinguish whether cello-oligosaccharides originate from decomposing lignocellulose (nutrient sources) or, instead, emanate from living and expanding plant tissue (virulence signals), and accordingly adapt its physiological response.

## Introduction

*Streptomyces* species are well-known soil-dwelling bacteria that actively participate in the recycling of organic matter mainly originating from plant residual biomass through diverse enzymatic systems (cellulases, amylases, xylanases,…) (1). Only few members of this genus have evolved from a saprophytic to a pathogenic lifestyle with *Streptomyces scabies* as model organism (2). Plant pathogenic *Streptomyces* are responsible for scab disease which can have severe consequences on production yields of economically important root and tuber vegetables (3). Thaxtomin A is the main virulence determinant produced by S. *scabies* which inhibits the cellulose biosynthesis process in underground expanding plant tissues (4-6). Cellobiose and cellotriose have been reported as the main carbohydrates triggering thaxtomin production and therefore the onset of the pathogenic behavior of S. *scabies* (7,8). Proteins of the cello-oligosaccharide-mediated induction of thaxtomin production consist of the CebEFG-MsiK ATP-binding cassette (ABC) transporter system (9, Figure 1). Control of the thaxtomin biosynthesis genes at the transcriptional level involves a double locking system with the cellulose utilization repressor CebR as the fastening key and the activator TxtR as the opening key (10,11). Despite the fact that the key players of the thaxtomin induction pathway have been recently characterized in detail, it is still unclear how S. *scabies* manages to sense cello-oligosaccharides as virulence signal and not as nutrient source. Here below we propose a series of features that could possibly permit *S*. *scabies* to avoid behaving as a pathogen if the conditions are more appropriate for a saprophytic lifestyle.

**Figure 1.**
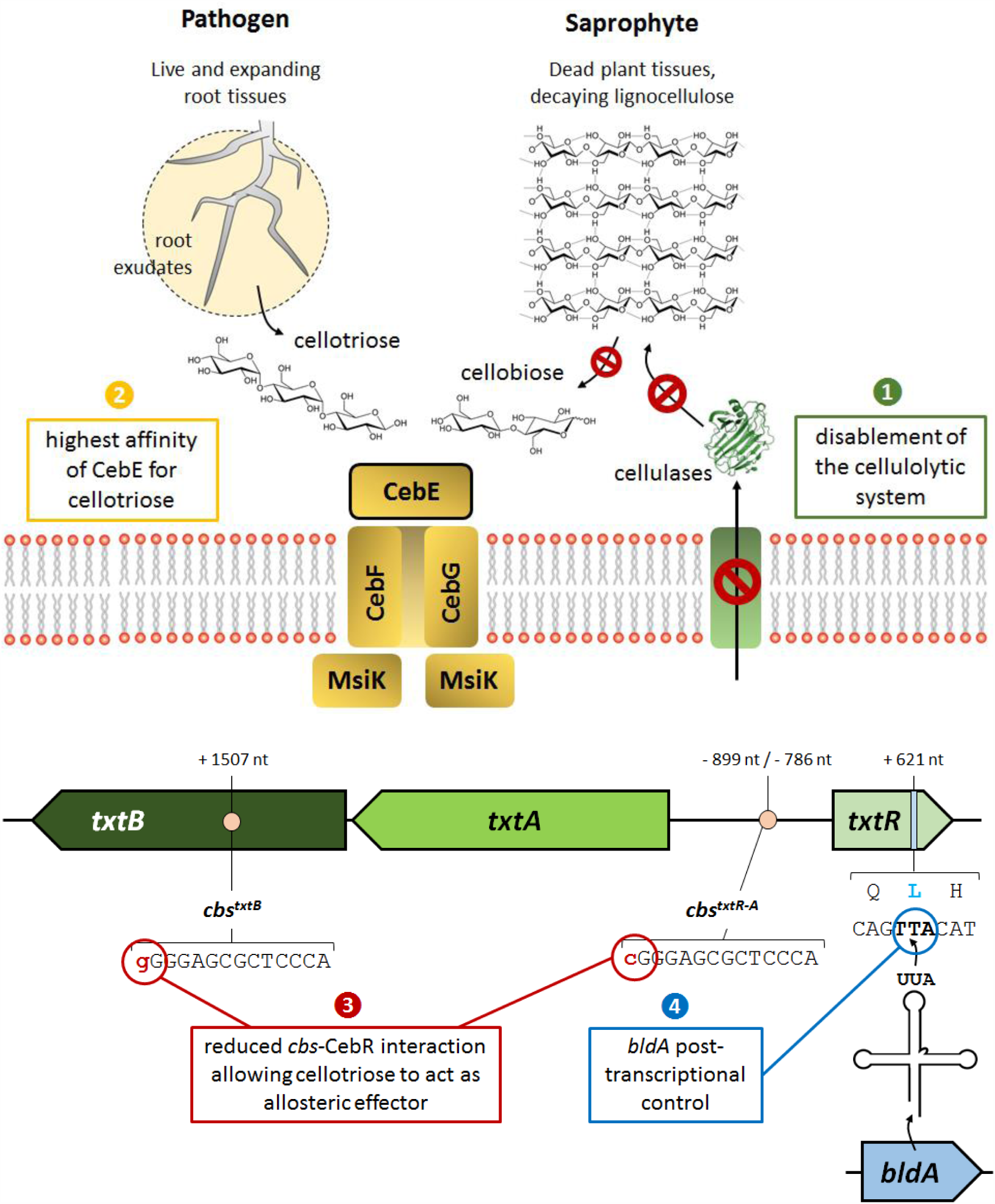
Genetic and physiological features predicted for adaptation of S. *scabies* to a pathogenic lifestyle built upon the perception of cello-oligosaccharides. 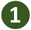 the disablement of the cellulolytic system; 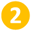 The increased affinity of the transporter sugar-binding CebE towards cellotriose (root exudates) instead of cellobiose (breakdown of cellulose); 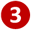 fine-tuned expression control of *txtR* by CebR; 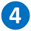 Post-transcriptional control of *txtR* by *bldA* encoding the leucyl-tRNA for the rare UUA codon.

## Disabling the cellulolytic system

Organisms able to actively participate in generating virulence elicitors are perceived as evolutionary more achieved compared to opportunistic pathogens that only passively wait for the looked-for signal to occur. However, when the induction of the pathogenicity system is mainly built upon sensing molecules originating from the most abundant polymer in organic soils (cellulose), the first step on the path to virulence should certainly differ from participating in the genesis of the signal. Having based its phytotoxin production on the use of cello-oligosaccharides is also very intriguing as the signal is first of all an important nutrient source that mostly originates from hydrolysis of perishing plant material. Therefore, it does not seem logic to use these cello-oligosaccharides as a trigger for virulence if releasing-hosts are dead. Therefore, a priority for S. *scabies* should entail avoiding the utilization of cellulose of decomposing plants if this species was to evolve to essentially behave as a pathogen. Indeed, S. *scabies* is unable to grow and use any type of cellulose when provided as the main nutrient source which prevents this organism to enzymatically generate their own virulence elicitors. How *S*. *scabies* disabled its complete extracellular enzymatic system dedicated to cellulose utilization is currently unknown and providing an answer to this question will most likely highlight a key mechanism on ‘how a saprophytic organism can become a plant pathogen’. The mutations involved must affect key proteins of induction and/or secretion mechanisms as S. *scabies* has a full set of genes of celluloclastic and cellulolytic systems *(cel* genes, Table 1) including enzymes belonging to the glycosyl hydrolase (GH) GH9, GH12, GH48 and GH74 families which are found only in *Streptomyces* with high cellulolytic activity (12). This paradox is reinforced by the abundance of perfect CebR-binding sites (cbs) found in S. *scabies* (14 *cbs* upstream of genes encoding putative cellulases, Table 1) as the same study indeed showed a positive correlation between the number of *cbs* found in different *Streptomyces* species and their ability to hydrolyze cellulose (12). Interestingly, when cellulose containing media are supplied with alternative carbon sources that do not impose carbon catabolite repression, a cellulase activity can be detected confirming that the cellulolytic system of S. *scabies* is functional (13; S. Jourdan, unpublished data). Thus, S. *scabies* needs to first use other carbon sources to initiate spore germination and early growth before considering cellulose as a possible nutritive substrate.

**Table 1.**
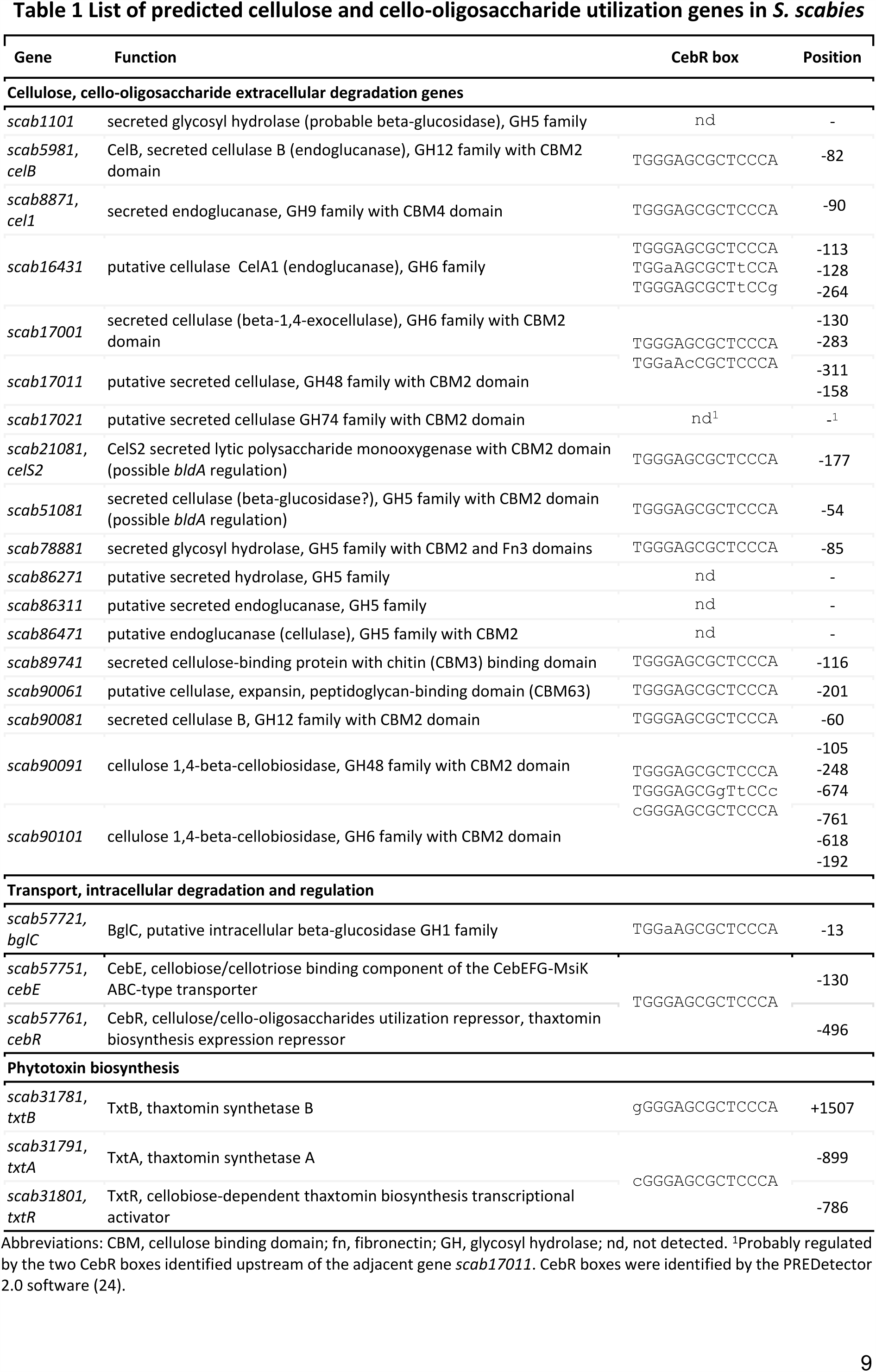
List of predicted cellulose and cello-oligosaccharide utilization genes in *S. scabies*

## How to distinguish identical molecules from live or dead plants?

The presence of cello-oligosaccharides in an environment devoid of potential living hosts could result from the degradation of lignocellulose derived from dead plant tissue by the action of cellulolytic microorganisms rather than by S. *scabies* itself as discussed above. Despite S. *scabies* containing cellulases that are transcriptionally and/or enzymatically silent, this strain possesses an efficient CebEFG-MsiK ABC-type transport system with a very high affinity (at nanomolar range) of CebE for cellobiose and cellotriose - respectively 100 to 1000 times higher than the affinity constants measured for the CebE of the highly cellulolytic species *Streptomyces reticuli* (14). This suggests that S. *scabies* would be adapted to behave as a commensal in soils with decaying plant cell walls. Nevertheless, S. *scabies* must have developed a mechanism which allows it to distinguish whether the source of cellobiose/cellotriose is derived from living or dead plant tissue in order to avoid counter-productive production of the phytotoxin thaxtomin. Although cellobiose, because of its reduced cost compared to cellotriose, is commonly used as an inducer of thaxtomin production in laboratory conditions, the trisaccharide has been reported to be a much better elicitor than the disaccharide (7). Interestingly, Johnson et al. showed that cellotriose (but not cellobiose) is present in the root exudates of growing radish seedlings as well as in tobacco cell suspensions (7). Instead, cellobiose is the main byproduct released by the action of the cellulolytic system while cellotriose is only marginally found from enzymatic degradation of cellulose. These observations suggest that cellotriose could be the signal molecule indicating the presence of a growing root network whereas cellobiose would signal dead plant cell walls to feed on. Expanding plant tissues releasing cellotriose not only constitute the site of invasion of the pathogen but also the site of action of thaxtomin. Johnson et al. also showed that the addition of thaxtomin resulted in an increase in the amount of cellotriose released by growing tissues. Thus, small amounts of cellotriose would be sufficient to initiate the production of thaxtomin which in turn would lead to the release of a greater amount of cellotriose which would trigger massive production of the phytotoxin. Thaxtomin itself, instead of the cellulolytic system, would be responsible to generate the inducer of its own biosynthesis as previously proposed by Johnson et al. This hypothesis implies that S. *scabies* would have evolved a mechanism able to monitor very low concentration of cellotriose emanating from root exudates. This seems to be the case as we have shown that the CebE lipoprotein of S. *scabies* has higher binding affinity for the trisaccharide (dissociation constant (Kd) for cellotriose ˜2 nM) than for the disaccharide (Kd cellobiose ˜14 nM) which is unusual for *Streptomyces* ABC-type sugar-binding components (15-17). Whether the presence of cellotriose and the absence of cellobiose is a constant in exudates from roots and tubers commonly damaged by S. *scabies* remains to be demonstrated. We are also currently investigating if the greater affinity of CebE for cellotriose instead of cellobiose is a specific feature of pathogenic *Streptomyces.* Site-directed mutagenesis will allow the identification of residues involved in this unusually high substrate affinity and bias towards the trisaccharide.

## Imperfect CebR cis-acting elements associated with virulence genes

The regulator CebR represses the expression of *cel* genes and the genes of the cello-oligosaccharide-mediated signaling cascade for thaxtomin production by binding to CebR-binding sites (cbs) with the 14-bp TGGGAGCGCTCCCA motif as the consensus palindromic sequence (10; Table 1). Genes that belong to a same regulon do not display the same expression level both under repressed or activated conditions. Indeed, for the proper response of a metabolic pathway, the expression of a certain category of genes has to be tightly inhibited while basal expression of others is required. In the case of the CebR regulon in S. *scabies*, one can imagine that genes involved in interpreting cello-oligosaccharides as a virulence signal might be differentially regulated than those strictly considering cellulose by-products as nutrient source. This differential expression pattern is in great part imposed by the distribution of one or more CebR cis-acting elements with different affinities in the gene’s upstream region (Table 1). From the 18 genes known or presumed to encode extracellular cellulases in S. *scabies*, 14 possess at least one perfect palindromic *cbs* in their upstream region (Table 1). Efficient activation of their transcription thus requires cellobiose, the best allosteric effector of CebR, while cellotriose only partially prevents its DNA-binding ability (10). The import of small amounts of cellotriose by the CebEFG-MsiK transporter would therefore have a much weaker effect on derepression of *cel* genes compared to the thaxtomin biosynthetic *(txtA* and *txtB)* and regulatory *(txtR)* genes which are more weakly bound by CebR due to the presence of imperfect cis-acting element (Figure 1). We therefore anticipate that cellotriose uptake from root exudates will first activate the expression of *txtR, txtA* and *txtB,* while cellobiose import from lignocellulose degradation will trigger expression of genes of the cellulolytic system.

## *bldA*-dependent developmental post-transcriptional control

The pathway-specific transcription factor TxtR is absolutely required for activation of the thaxtomin biosynthetic genes (7). This essential gene for the induction of the S. *scabies* pathogenicity is not only controlled by CebR at the transcriptional level but also possesses a TTA codon at nucleotide position 621 (18; Figure 1); TTA codons are extremely rare in GC-rich *Streptomyces* species. The UUA codon in *Streptomyces* mRNA can only be translated into a leucine by the leucyl-tRNA encoded by the *bldA* gene. Post-transcriptional regulation by *bldA* is a common feature associated with physiological and morphological differentiation processes in all *Streptomyces* (19,20). The inactivation of *bldA* results in a so-called bald (non sporulating) phenotype with concomitant impaired production of their specialized metabolites (antibiotics, siderophores, …). Induction of *bldA* transcription occurs in order to allow the translation of mRNA encoding proteins that enable adaptation to transient stress conditions and/or occur during late exponential growth, that is, when *Streptomyces* switch from primary to secondary metabolism and trigger their developmental processes (21). This additional post-transcriptional control mediated by *bldA* would prevent S. *scabies* from producing thaxtomin even in the presence of cello-oligosaccharides unless other factors (stress, population density, life cycle stage,…) affecting *bldA* expression are present, indicating the proper timing to switch from the saprophytic to the pathogenic lifestyle.

## Concluding remarks and perspectives

Previous reviews have listed and discussed in detail the acquired large genetic material - at the gene or gene cluster level - for S. *scabies* to behave as a plant pathogen (22,23). Although these major virulence determinants are now easier to identify as we entered the era of high-speed and cost-efficient genome sequencing, finding the discrete but equally essential genetic changes necessary for transition from saprophytic to pathogenic lifestyles remains a challenging task. Our search for the subtle genetic causes that would possibly explain the physiological adaptation of S. *scabies* includes mutations that (i) modify the affinity of the CebE sensor for the most appropriate virulence cue cellotriose, (ii) weaken the interaction strength of the transcriptional repressor CebR for its *cis*-acting elements, and (iii) impose post-transcriptional control by the use of the rare TTA codon ensuring that toxin production correlates with the developmental transition. The mutations associated with the disablement of the cellulolytic system remain enigmatic in the current state of our knowledge but might include some of those cited above. For instance *scab51081* and *scab21081 (celS2)* are possible targets of BldA post-transcriptional regulation as they have a TTA codon at nucleotide positions 13 and 22, respectively. In addition, tight CebR mediated repression is also suggested on the expression of *scab16431, scab17001, scab17011, scab90091,* and *scab90101* as evidenced by multiple CebR boxes in their upstream region (Table 1). The hypotheses presented and discussed here require further experimental validation and are probably only a part of all minor changes that appeared in the ancestors of pathogenic *Streptomyces* species once they acquired the thaxtomin biosynthesis cluster.

## Acknowledgments

S.J.’s work is supported by an Aspirant grant from the FNRS (Grant R.FNRS.2898-4-F; 1.A250.13). S.R. is a FRS-FNRS research associate. This work is supported in part by the Belgian program of Interuniversity Attraction Poles initiated by the Federal Office for Scientific Technical and Cultural Affairs (PAI no. P7/44) and by the FNRS (research project R.FNRS.3342; T.0006.14-PDR).

